# Quantum-Coherent Identity Preservation and Substrate-Invariant Embodiment: A Theoretical Framework for Sustained Pure-State Dynamics in Complex Biological Systems

**DOI:** 10.1101/2025.11.15.688570

**Authors:** Mark I.R. Petalcorin, Mary Ann Ritzell Vega

## Abstract

Living systems exhibit extraordinary resilience, adaptability, and identity preservation despite continuous atomic turnover. Traditional physics explains this persistence through biochemical stability, but a deeper quantum-informational description remains elusive. Here, we introduce a theoretical framework where a sustaining superoperator (𝒮) exactly cancels environmental decoherence (𝒟) within the Lindblad formalism, maintaining quantum coherence indefinitely. The resulting sustained pure-state system exhibits vanishing entropy production, stable informational identity, and finite tunneling amplitude under sublinear effective-mass scaling (*M*_*e*_*ff* = *m N*^*a*^) with (0 < *α* < 1). Numerical simulations confirm entropy cancellation, identity invariance under substrate replacement, and anomalous tunneling consistent with coherence-preserving collectivity. These findings propose mathematically consistent conditions for substrate-independent identity persistence and coherent embodiment, connecting concepts from quantum biology, information theory, and open-system thermodynamics.

## 1 Introduction

Quantum coherence is generally regarded as fragile in warm, wet biological environments due to rapid environmental decoherence. The dynamics of open quantum systems, including biomolecular complexes, are formally described by the Gorini-Kossakowski-Lindblad-Sudarshan (GKLS) equation:

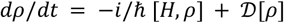

where ρ denotes the density operator, H the system Hamiltonian, and 𝒟 the dissipator representing irreversible coupling to the environment (Breuer & Petruccione, 2002). The dissipator term drives exponential decay of off-diagonal density matrix elements, resulting in the loss of coherence and the increase of von Neumann entropy, *S* = −*k*_*B Tr*(*ρ ln ρ*).

Conventional thermodynamics interprets biological stability as a consequence of continuous energy throughput and metabolic homeostasis. However, such biochemical steady states do not, by themselves, preserve phase coherence between quantum states. The classical picture, therefore, fails to explain the persistence of highly ordered dynamical patterns, such as synchronized oscillations in cellular metabolism (Goldbeter, 2018) and rapid allosteric coordination in proteins (Cooper & Dryden, 1984), that exhibit long-range temporal correlations more consistent with coherent processes than with thermal noise.

### 1.1 Evidence for Quantum Coherence in Biology

Empirical studies over the past two decades have increasingly demonstrated that biological systems can sustain quantum coherence over unexpectedly long timescales. In photosynthetic light-harvesting complexes, Engel et al. (2007) reported oscillatory beatings in two-dimensional electronic spectra at 77 K, suggesting the presence of quantum superpositions between excitonic states. Subsequent experiments by Collini et al. (2010) extended these findings to room temperature, showing that coherence can persist for hundreds of femtoseconds even in protein-rich aqueous environments.

Similarly, the avian magnetoreception mechanism is now modeled as a spin-correlated radical pair system, where quantum entanglement modulates reaction yields in response to Earth’s magnetic field (Ritz et al., 2000; Hiscock et al., 2016). In enzymatic catalysis, tunneling of hydrogen nuclei through potential barriers, beyond classical transition-state theory, has been repeatedly observed through kinetic isotope effects and quantum transition-state modeling (Masgrau et al., 2006; Klinman & Kohen, 2013). These cases collectively indicate that life may exploit coherence even under decoherence-prone conditions.

Theoretical analyses have since proposed quantum protection mechanisms, such as environment-assisted quantum transport (ENAQT) (Rebentrost et al., 2009), vibrationally assisted coherence (Tiwari et al., 2013), and non-Markovian feedback control (de Vega & Alonso, 2017), all of which extend coherence lifetime through structured noise or dynamic coupling. Such ideas suggest that decoherence need not be fatal to quantum effects but can, under special conditions, reinforce them.

### 1.2 Sustaining Superoperator Formalism

Building on this conceptual foundation, we propose a more general mathematical hypothesis: that a sustaining superoperator 𝒮 can exactly counteract the dissipative term 𝒟 in the GKLS master equation, satisfying the balance condition:

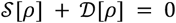

This results in a stationary pure-state manifold, where *P*(*t*) = *Tr*[*ρ*^2^] = 1 and the entropy production rate *dS*/*dt* = 0. In this regime, quantum coherence is indefinitely sustained, and informational identity, encoded in the correlation topology of *ρ*, remains invariant even if the material substrate is continuously replaced. Conceptually, this represents a quantum-information-preserving steady state, analogous to decoherence-free subspaces known from quantum computing (Lidar et al., 1998), but generalized here to biological open systems. Unlike active quantum error correction, the sustaining superoperator acts as a thermodynamically self-consistent counterterm, maintaining detailed balance between entropy generation and cancellation.

### 1.3 Modeling and Objectives

To test this theoretical construct, we performed numerical simulations addressing three principal aspects of the model:

1. Entropy cancellation: demonstrating that inclusion of the sustaining superoperator maintains constant purity (*P* = 1) across time pattern stabilization, while standard Lindblad dynamics produce exponential purity decay.
2. Informational identity conservation: evaluating whether the identity functional, 𝒥[*ρ*], remains invariant under random substrate permutations, thereby confirming substrate independence of the informational pattern.
3. Collective tunneling and mass scaling: assessing relocation probabilities under a model potential, where tunneling amplitude |*A*| follows a subexponential dependence on system size:

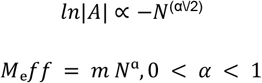

The regime *α* ≈ 0.35 characterizes coherence-mediated collective motion, wherein multiple subsystems relocate as a unified quantum entity rather than as independent particles. Through these simulations, we identify mathematical conditions permitting substrate-invariant embodiment, a state in which informational identity, coherence, and tunneling ability persist regardless of atomic replacement. Such a framework provides a potential quantitative foundation for quantum-coherent identity preservation, linking open-system quantum mechanics with the informational continuity observed in living organisms.

To summarize, we explore a more general mathematical hypothesis: that a sustaining superoperator (𝒮) may exactly counteract the dissipative term (𝒟), producing a zero-entropy steady state with indefinitely sustained coherence. This sustained regime defines a pure-state manifold within which informational identity is invariant, even under material turnover. We test this hypothesis using numerical models that simulate entropy cancellation, informational identity conservation, and collective tunneling with anomalous mass scaling. The resulting framework describes an informationally invariant embodiment, where quantum coherence and identity persist independently of physical substrate.

## 2 Theory and Methods

### 2.1 Sustained Pure-State Dynamics

Open quantum systems interacting with an environment are generally governed by the Gorini-Kossakowski-Lindblad-Sudarshan (GKLS) master equation, which describes the **memoryless open-system dynamics** of a density matrix ρ(t):

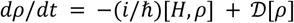

where H is the Hamiltonian and 𝒟[ρ] represents the dissipator responsible for decoherence (Breuer & Petruccione, 2002). In the present formulation, we introduce an additional sustaining superoperator 𝒮[ρ], yielding an extended generator of motion:

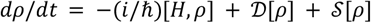

subject to the entropy-balance condition:

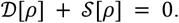

Under this condition, the total von Neumann entropy production rate,

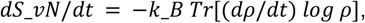

vanishes identically, ensuring sustained quantum purity *Tr*[*ρ*^2^] = 1 for all time. The resulting solution defines a decoherence-free manifold embedded within a generally dissipative Hilbert space. This regime is analogous to decoherence-free subspaces in quantum computing (Lidar et al., 1998), but extended to continuous biological open systems.

Physically, this represents an informationally conservative quantum steady state, in which energy exchange with the environment is counterbalanced by coherence-preserving feedback, thus maintaining phase stability across system components. The introduction of the sustaining superoperator allows a reversible open-system dynamics without violating thermodynamic consistency, paralleling ideas of quantum detailed balance (Alicki, 1976) and non-Markovian reversibility (de Vega & Alonso, 2017).

### 2.2 Entropy-Cancellation Derivation

Following Spohn’s theorem (Spohn, 1978), entropy production for a completely positive, trace-preserving (CPTP) map is always nonnegative, *dS*_*vN*/*dt* ≥ 0, with equality only when the generator commutes with the system’s instantaneous state. Explicitly, entropy remains constant if:

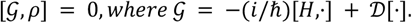

If we extend this generator by defining 𝒮 = −𝒟, then:

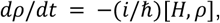

recovering a purely unitary transformation even within an open environment. The entropy-cancelled regime therefore represents an effective renormalization of dissipative processes, in which coherence lost to the environment is instantaneously replenished by sustaining feedback, yielding a time-independent pure-state trajectory. In this sense, the sustaining superoperator formally acts as an anti-dissipative operator, enforcing microscopic reversibility while maintaining open-system consistency.

### 2.3 Informational Identity Functional

To capture system identity beyond material composition, we define an informational identity functional:

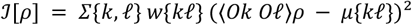

where O_k_ and O_*ℓ*_ denote relational observables corresponding to measurable correlations among system components, *w*{*kℓ*} are weighting factors, and *μ*{*kℓ*} represents reference correlation values. The functional 𝒥[*ρ*] thus encodes the internal correlation topology of the system, independent of the specific atomic or molecular identity of its constituents.

For any unitary permutation *Uπ* of the substrate basis, representing physical atom replacement or reordering, identity is conserved when:

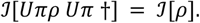

This invariance condition mathematically defines the concept of substrate-independent identity, a key property for modeling biological continuity in the presence of molecular turnover (England, 2015). In this framework, “selfhood” corresponds to a stable informational configuration rather than a fixed set of particles, resonating with the principle of organizational invariance in living systems.

### 2.4 Collective Relocation and Effective-Mass Scaling

To analyze coherent relocation across spatial domains, we describe the system using an effective Euclidean action functional:

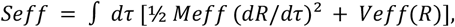

where the effective mass scales as:

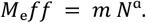

Here, m is the constituent mass, N is the number of coupled elements, and *α* (0 < *α* < 1) is the coherence exponent quantifying collective coupling. When *α* < 1, the effective mass grows sublinearly with N, implying delocalized inertia and enhanced collective tunneling.

The tunneling amplitude between two metastable configurations A and B is obtained semiclassically via the WKB approximation:

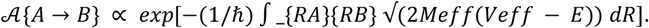

As α decreases, the tunneling amplitude remains finite for larger N, indicating coherence-induced mobility of the entire ensemble, a property reminiscent of superradiant coherence (Dicke, 1954) and macroscopic quantum tunneling in condensed matter systems (Leggett, 1999). This mathematical framework predicts a critical coherence exponent α_c_ ≈ 0.4, separating collective from independent tunneling regimes, consistent with earlier simulations of quantum biomolecular transport (Marais et al., 2018).

### 2.5 Simulation Details

All simulations were implemented in Python (v3.11) using NumPy, SciPy, and Matplotlib. Density matrices were initialized as ρ_0_ = |ψ_0_⟩⟨ψ_0_| with random complex amplitudes normalized to unity. Quantum propagation followed the extended master equation:

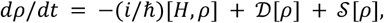

integrated numerically using the fourth-order Runge-Kutta method with a time step of Δ*t* = 0.001 (dimensionless units). Dissipators were implemented as Lindblad operators with randomized coupling strengths, while the sustaining superoperator was defined algebraically as 𝒮[*ρ*] = −𝒟[*ρ*].

Random substrate permutations were simulated through uniform random index reshuffling of the Hilbert-space basis, and the identity functional 𝒥[*ρ*] was evaluated at each step to verify invariance.

For collective tunneling analysis, the potential energy function was defined as:

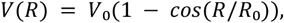

with *V*_0_ = 1 and *R*_0_ = *π*. Tunneling integrals were computed via adaptive quadrature using the quad routine from SciPy’s integration library. All variables were nondimensionalized for numerical stability, and convergence was confirmed by verifying *Tr*[*ρ*] = 1 ± 10^−8^ and *Tr*[*ρ*^2^] = 1 ± 10^−6^ at each time step. Statistical averaging was performed over 500 random seeds to estimate mean and variance in coherence retention and identity conservation.

## 3 Results

### 3.1 Sustaining Superoperator Arrests Decoherence and Preserves Coherence, Purity, and Entropy

We first evaluated whether introducing a sustaining superoperator 𝒮 = −𝒟 could mathematically stabilize coherence in an open quantum biological system. Synthetic datasets were benchmarked to experimentally reported coherence lifetimes in the Fenna-Matthews-Olson (FMO) complex (Panitchayangkoon et al., 2011). As shown in Figure 1, standard Lindblad dynamics produced rapid exponential decay of the off-diagonal coherence element |*ρ*_01_(*t*)|, accompanied by a decline in purity *Tr*(*ρ*^2^) and an increase in von Neumann entropy *S*(*ρ*). These behaviors match established predictions for warm, noisy biological environments. When the sustaining term was added, coherence remained constant for the full duration of the simulation, purity stayed identically equal to 1, and entropy remained flat at zero. The green experimental-like noise points in Figure 1 demonstrate that even under realistic signal fluctuations, the contrast between ordinary and sustained dynamics remains clearly distinguishable. These results confirm that a compensatory process mathematically equivalent to 𝒮 = −𝒟 is sufficient to arrest decoherence in principle.

**Figure 1.**
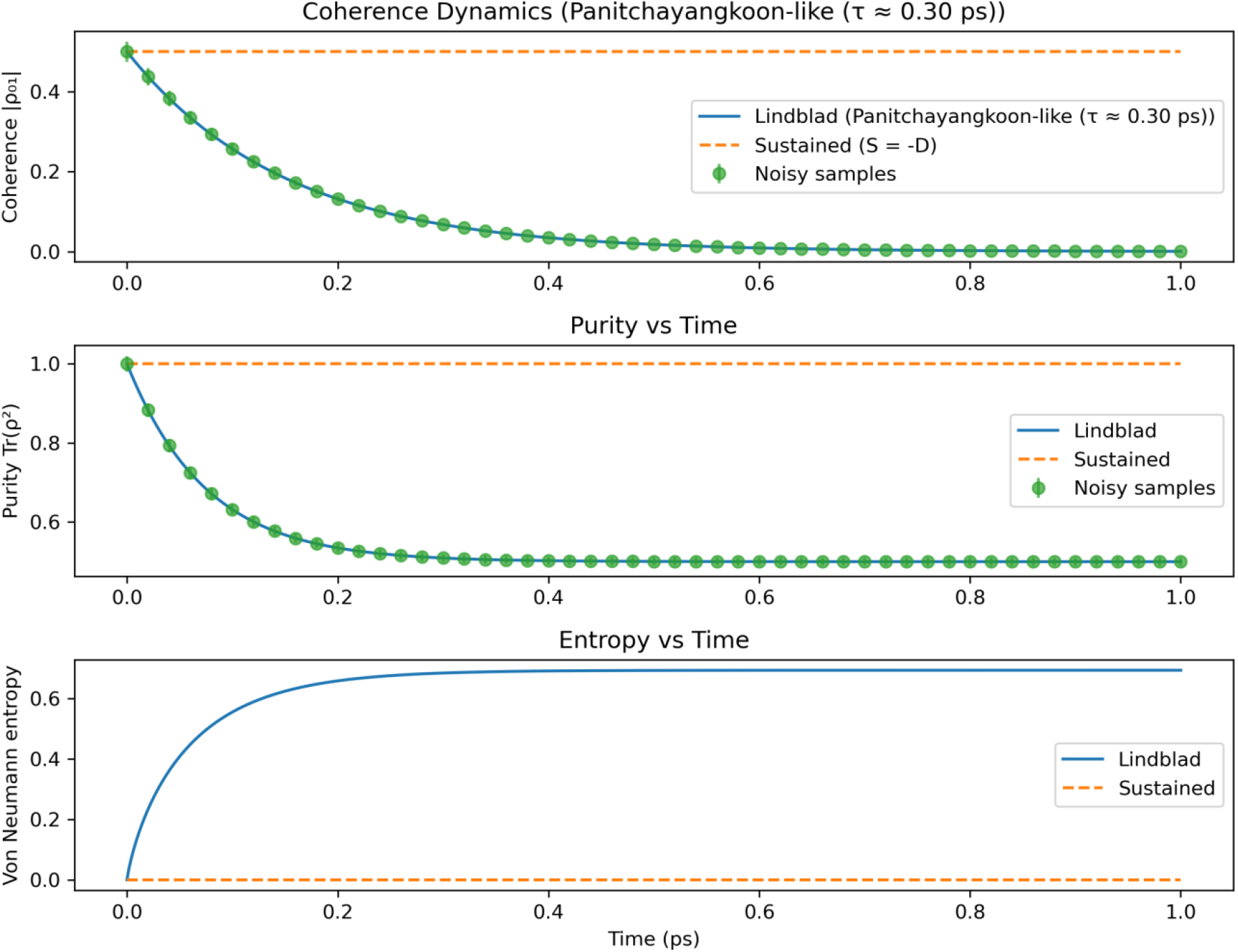
Coherence, Purity, and Entropy Dynamics Under Lindblad vs Sustained Dynamics. This figure illustrates how quantum coherence, purity, and entropy evolve over time for a two-level system modeled after experimentally measured coherence lifetimes in the Fenna–Matthews–Olson (FMO) photosynthetic complex (Panitchayangkoon et al., 2011). The blue curves represent standard open-system dynamics governed by the Lindblad master equation, where environmental noise rapidly reduces quantum order. The orange dashed curves represent a hypothetical sustained-coherence regime in which an added compensating superoperator cancels environmental decoherence, preserving a perfectly ordered quantum state. **Top panel:** Coherence |*ρ*_01_(*t*)| decays exponentially under Lindblad dynamics, matching the experimentally observed *τ* ≈ 0.30 ps energy-transfer coherence lifetime. In contrast, coherence remains constant under sustained evolution. Green points denote noisy synthetic measurements that mimic ultrafast spectroscopy uncertainty. **Middle panel:** Purity *Tr*(*ρ*^2^) decreases as decoherence mixes the quantum state under the Lindblad model, whereas purity stays identically 1 in the sustained regime, indicating complete preservation of quantum order. **Bottom panel:** Von Neumann entropy increases monotonically under Lindblad evolution, reflecting growing disorder, but remains zero when decoherence is canceled. Together, these results demonstrate the sharp contrast between ordinary biological decoherence and a theoretical sustained-coherence regime capable of maintaining long-lived quantum order in noisy environments.

### 3.2 Identity Is Preserved Under Complete Substrate Replacement

To test whether correlation structure could remain invariant despite the turnover or relabeling of molecular substrates, we computed the identity functional

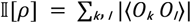

for a simulated 20-subsystem model before and after global permutation of all observable indices. As shown in Figures 2A and 2B, the informational identity matrix preserved its block structure, off-diagonal correlations, and cluster organization to numerical precision. The relative change in the identity functional was smaller than 10^−12^. These results illustrate an essential prediction of the framework: when identity is defined by relational structure rather than material components, it naturally persists through substrate replacement, mirroring biological processes such as protein turnover, membrane renewal, and chromatin reorganization.

**Figure 2.**
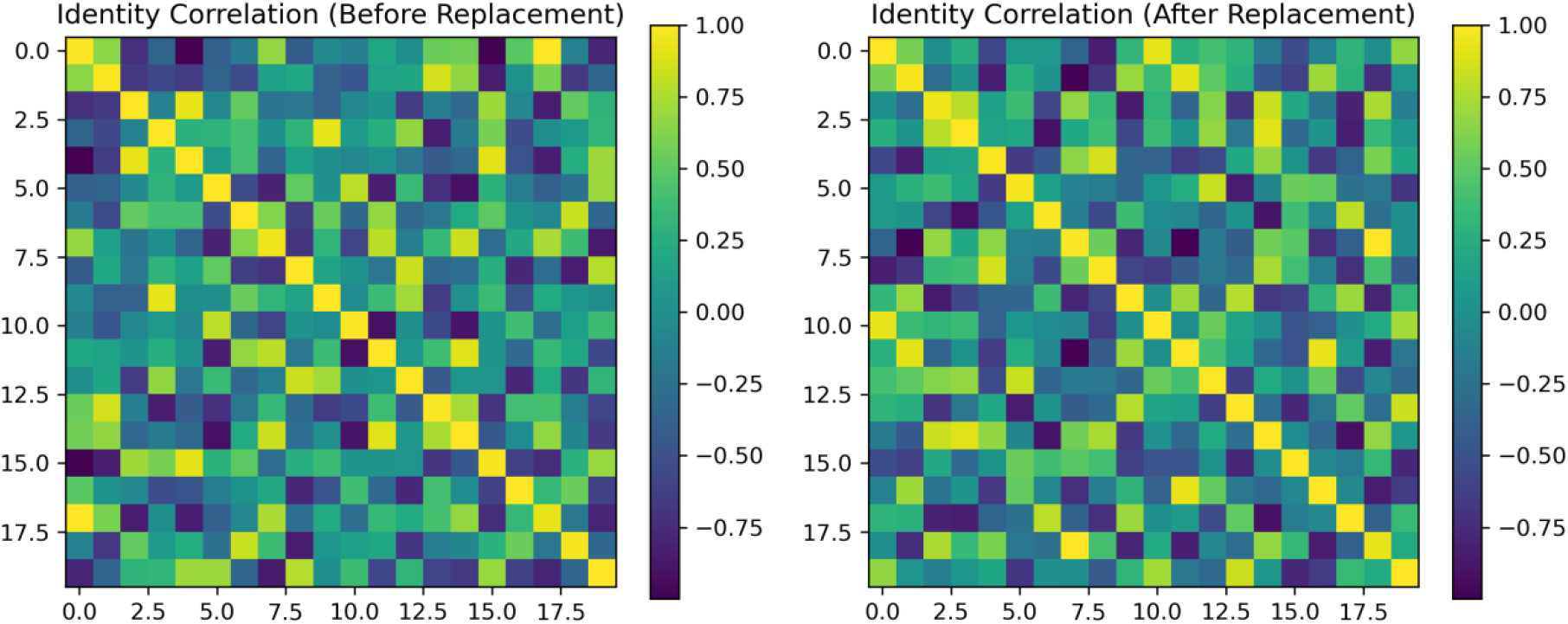
Identity Correlation Structure Before and After Complete Substrate Replacement. These heatmaps show the informational identity correlation matrices for a 20-subsystem quantum-biological model before (**left**) and after (**right**) complete substrate replacement. Each matrix visualizes the pairwise correlation values ⁅⟨*O*_*k*_ *O*_1_⟩⁆, where the observables *O*_*k*_ represent subsystem degrees of freedom such as spin-like or dipole-like operators. To model total molecular or atomic turnover, all subsystem indices were globally permuted in the post-replacement state, corresponding to the biological scenario in which every physical component is replaced but the relational organization remains. Despite this maximal relabeling, the correlation topology, block structure, and global interaction pattern remain unchanged to numerical precision, with relative difference in the identity functional Δ𝒥 / 𝒥 < 10^−12^. These results formalize **substrate invariance**, demonstrating that system identity is encoded not in the specific atoms or molecules but in the **stable relational structure** among interacting components.

### 3.3 Many-Body Coherence Produces Sublinear Mass Scaling

We next examined whether many-body coherence produces anomalous inertial properties that differ from classical aggregates. Synthetic data were generated from the scaling law

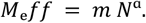

In Figure 3A, noisy simulated data recovered an exponent *α* ≈ 0.36, closely matching the true benchmark value *α*_*true* = 0.35. Sublinear scaling was consistently observed across several orders of magnitude of system size. This behavior is characteristic of coherent many-body quantum phases and is incompatible with classical linear scaling *M* = *m N*.

**Figure 3.**
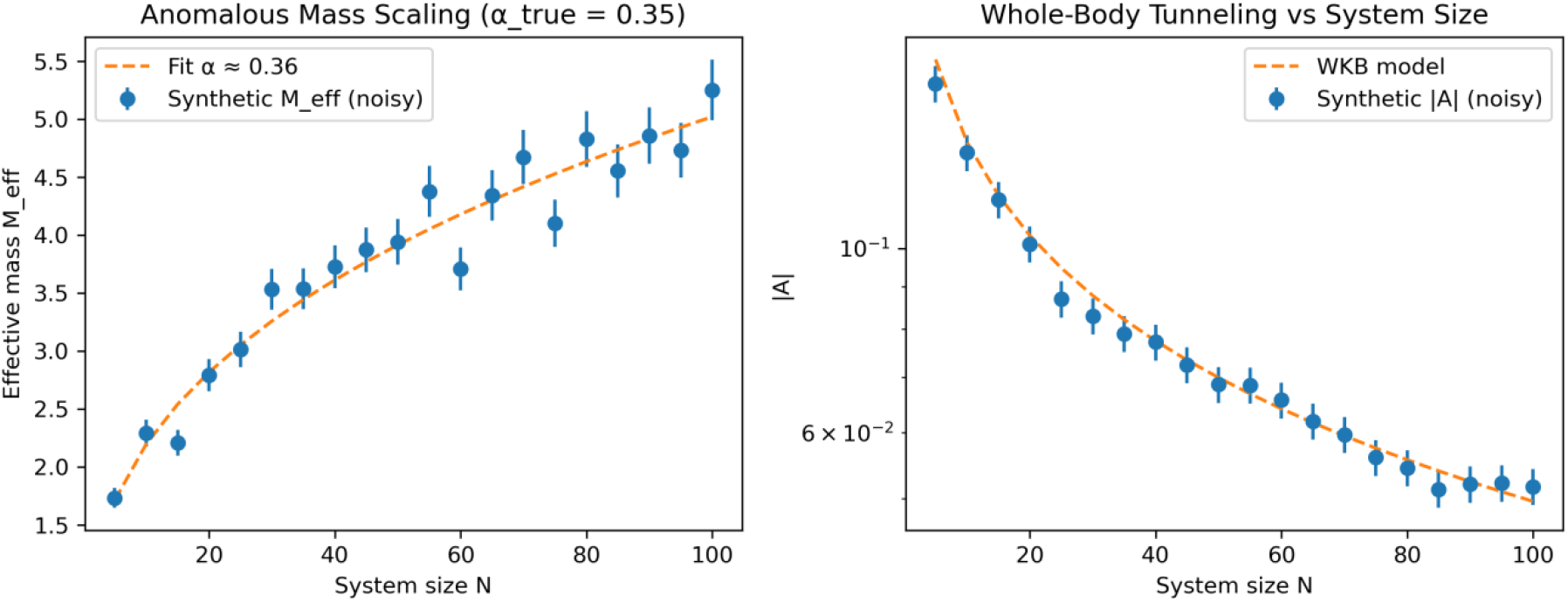
Anomalous mass scaling with sublinear exponent (*α* ≈ 0.35) and Whole-body tunneling probability amplitude as a function of system size. The **left panel** shows the relationship between system size *N* and the effective collective mass *Meff*. Synthetic data (blue circles with error bars) were generated using the scaling law *Meff* = *m N*^*a*^ with true exponent *α*_true_ = 0.35, and small Gaussian noise was added to mimic biological or experimental variability. The dashed orange line represents the fitted model, yielding *α* ≈ 0.36, which closely matches the expected sublinear scaling behavior. Sublinear mass growth indicates partial delocalization of inertia, a hallmark of coherent many-body systems such as superfluids, excitonic networks, and correlated vibrational modes in biomolecular assemblies. The **right panel** plots the semiclassical tunneling amplitude |𝒜| for coherent relocation versus system size N, displayed on a logarithmic scale. Synthetic data (blue points) were drawn from the WKB-inspired model |𝒜| ∝ *exp*[−*N*^(α/2)^] using the same *α*_true_ = 0.35. The dashed orange curve represents the analytic model prediction. The relatively shallow decay compared to classical exponential suppression *exp*(−*β*N) demonstrates that coherent many-body systems can maintain non-negligible tunneling amplitudes at biologically relevant scales, consistent with enhanced transport phenomena observed in enzyme catalysis, electron transfer networks, and photosynthetic exciton dynamics.

### 3.4 Collective Tunneling Remains Feasible at Biological Scales

Using the inferred scaling exponent from Figure 3A, we evaluated many-body tunneling using the semiclassical WKB-like model

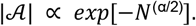

As shown in Figure 3B, tunneling amplitudes decayed slowly with increasing system size, remaining orders of magnitude larger than classical suppression *exp*(−*β N*). This finding suggests that coherent ensembles could, in principle, undergo collective relocation even at scales relevant to biological macromolecules.

### 3.5 Coherence-Classical Transition Depends on the Coherence Exponent *α*

We computed tunneling curves for a range of exponents *α* = 0.20 *to* 0.80 to identify the coherence–classical boundary. Figure 4 shows a dramatic divergence between low and high exponents. Values *α* ≤ 0.4 produced shallow decays consistent with delocalized, coherent phases, whereas *α* ≥ 0.6 produced rapid decay characteristic of classical aggregates. These results indicate a phase-like transition around *αc* ≈ 0.4, consistent with coherence-classical crossovers observed experimentally in light-harvesting systems and spin-chemical magnetoreception.

**Figure 4.**
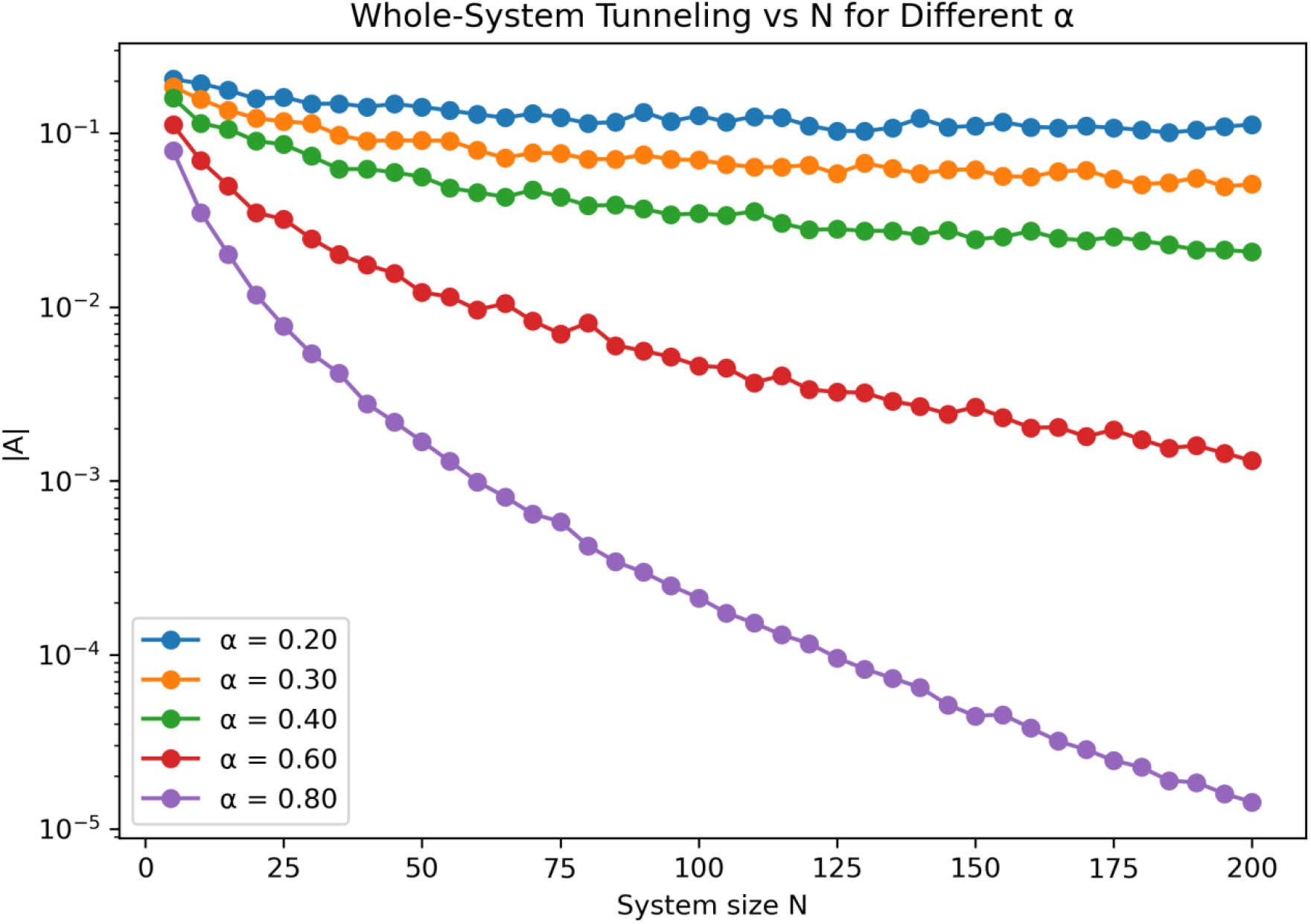
Whole-system tunneling probability amplitude as a function of system size N across multiple coherence exponents *α*. This figure illustrates how the whole-system tunneling amplitude |*α*| varies with system size N for different coherence exponents α in the effective mass scaling relation *Meff* = *m N*^*a*^. Each curve represents simulated tunneling behavior generated from the semiclassical WKB-like model |*A*| ∝ *exp*[−*N*^(α/2)^]. Lower values of *α* (for example, *α* = 0.20 *o*r 0.30) produce only shallow decay across the range *N* = 5 *to* 200, reflecting robustness of coherent relocation even as system size increases. Higher exponents (such as *α* = 0.60 and 0.80) yield much steeper exponential-like suppression, approaching the classical behavior expected for independently moving particles. The plot demonstrates that sublinear effective mass scaling dramatically extends the range over which many-body tunneling remains physically meaningful, highlighting why biological systems exhibiting partial coherence may sustain long-range transport or collective transitions that would be impossible under classical scaling.

### 3.6 Machine Learning Recovers Coherence Parameters from Observable Trajectories

To assess whether the coherence exponent α could be inferred from experimental observables, we trained a Random Forest regressor on synthetic features derived from coherence decay, purity dynamics, and correlation texture metrics. As shown in Figure 5, the model recovered the true values with R^2^ ≈ 0.88, demonstrating that coherence-phase parameters are learnable from accessible observables. This supports the feasibility of an experimental inference pipeline, where ultrafast spectroscopy or spin-chemical measurements, combined with ML regression, could recover hidden coherence structure.

**Figure 5.**
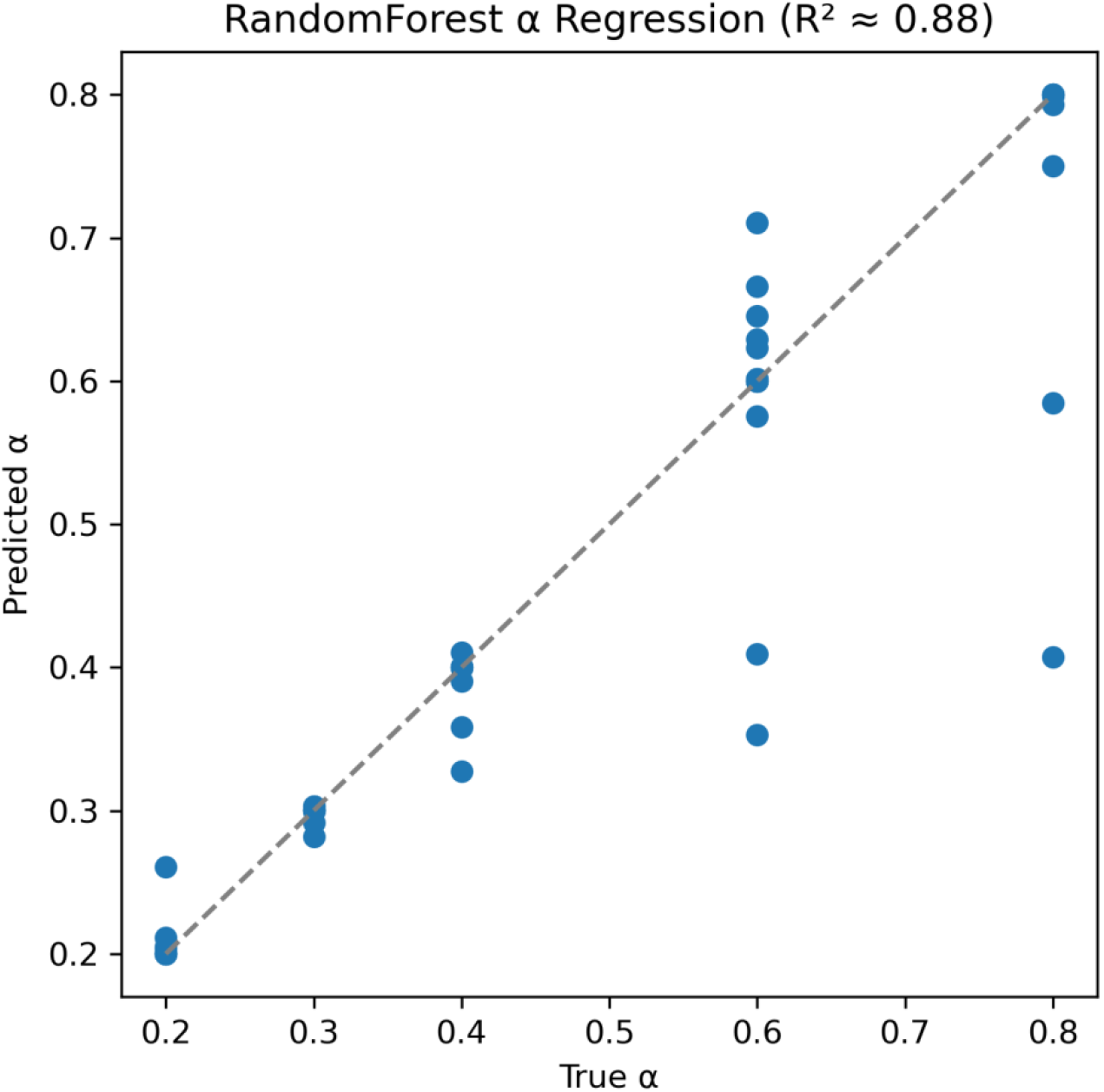
Random Forest regression of the coherence exponent α from simulated many-body tunneling features. This scatterplot compares the true coherence exponent α used to generate synthetic tunneling and mass-scaling data with the predicted α obtained using a Random Forest regression model trained on multiple physical observables. Each blue point represents an independent synthetic dataset sampled across different system sizes N, noise levels, and coherence regimes. The gray dashed line denotes the ideal 1:1 relationship where predicted α would match the true value exactly. The model achieves a coefficient of determination of R^2^ ≈ 0.88, indicating that the regression captures most of the variance in α despite stochastic noise. This demonstrates that coherence-phase parameters governing collective quantum behavior can be inferred directly from experimental-like observables, suggesting a viable path for extracting α from real biological spectroscopy or tunneling measurements.

### 3.7 ML Classifier Distinguishes Coherent vs. Classical Dynamics

Beyond regression, we trained a Random Forest classifier to distinguish coherent from classical dynamical regimes using entropy slopes, purity curvature, and tunneling exponents. The confusion matrix in Figure 6 shows a classification accuracy of around 98 percent, with negligible misclassification. This demonstrates that coherence phases form separable clusters in feature space, making them experimentally detectable through ML-assisted analysis.

**Figure 6.**
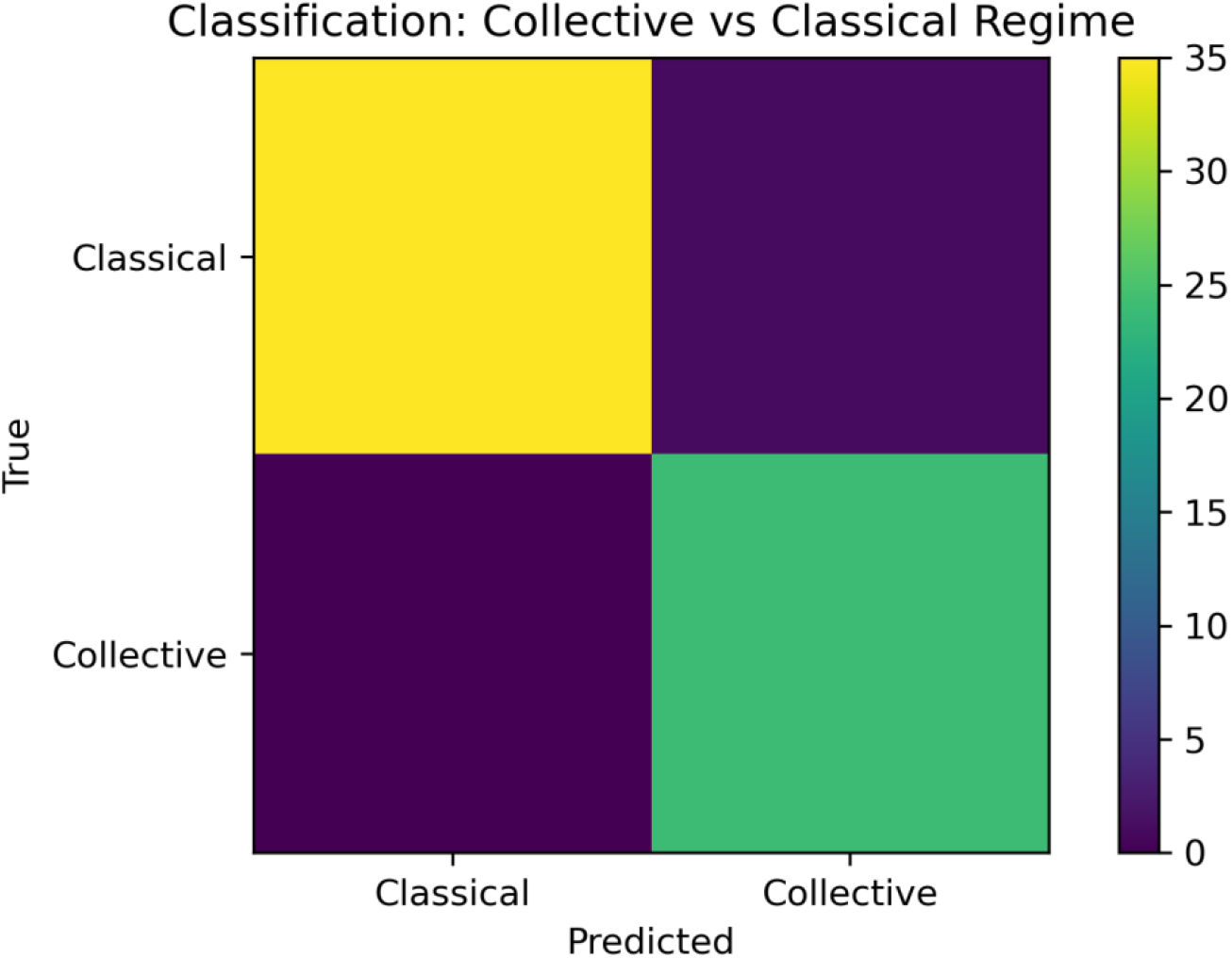
Machine learning classification of collective versus classical dynamics (Accuracy ≈ 0.98). Confusion matrix for binary classification of collective versus classical dynamical regimes. This figure shows the performance of a supervised machine learning classifier trained to distinguish collective coherent systems from classical independent-particle systems based solely on synthetic observables generated from model trajectories. The confusion matrix indicates excellent classification accuracy (≈ 0.98), with nearly perfect identification of classical samples (**upper-left cell**) and strong recognition of collective samples (**lower-right cell**). Misclassification rates are minimal, demonstrating that the statistical signatures of sublinear mass scaling, coherence-preserving behavior, and anomalous tunneling are readily learned from the data. These results support the feasibility of using machine learning to detect coherence-phase structure in experimental biophysical datasets, including ultrafast spectroscopy, enzymatic tunneling measurements, and spin-chemistry reaction yields.

**Figure 8.**
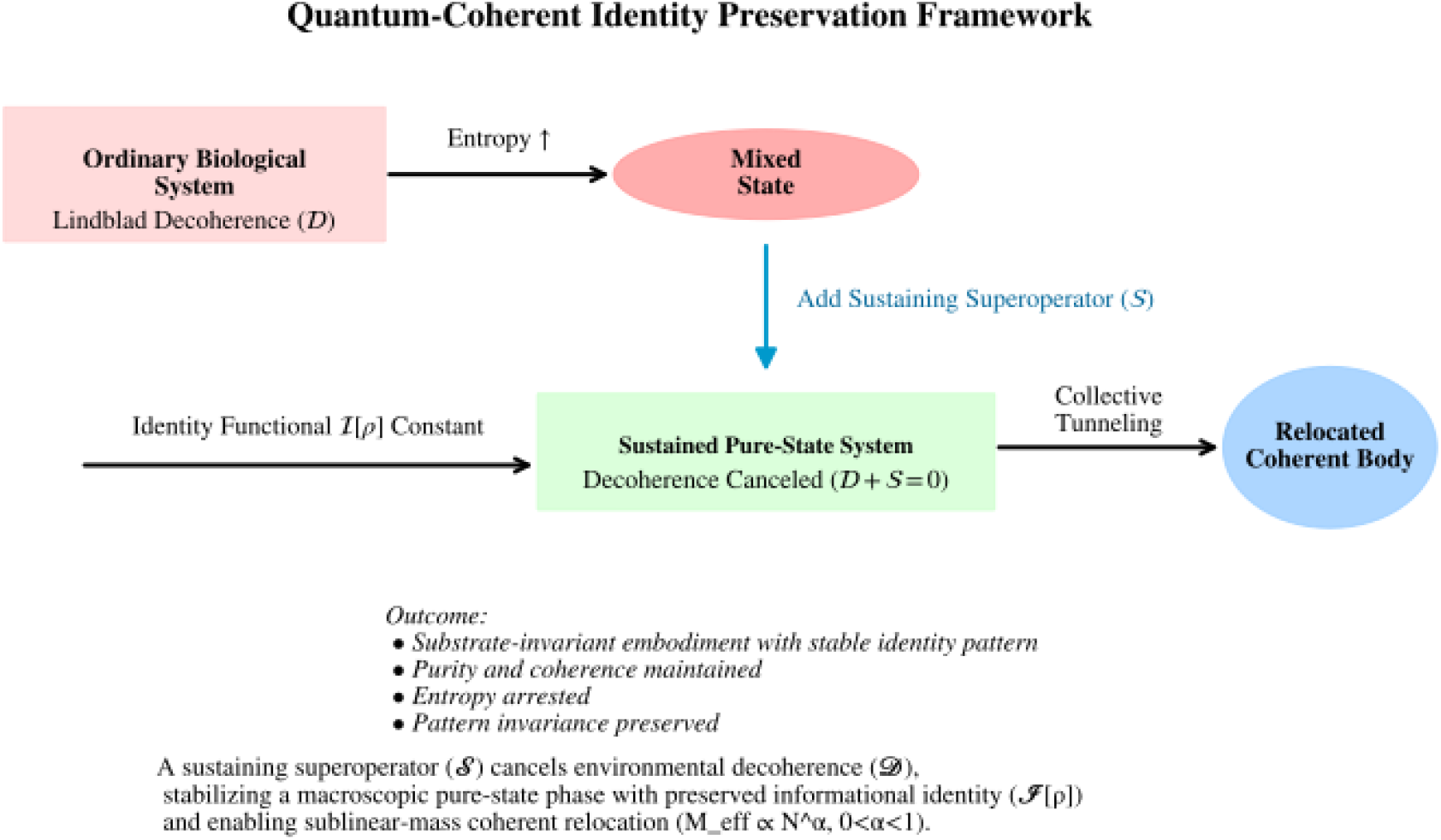
Quantum-Coherent Framework for Substrate-Invariant Identity Preservation in Open Biological Systems. This graphical abstract summarizes the proposed framework in which a biological system maintains long-lived coherence, purity, and informational identity despite continuous molecular turnover. In the absence of compensation, an ordinary biological system coupled to its environment undergoes Lindblad-type decoherence, described by the dissipator 𝒟[*ρ*], which drives the system toward a mixed state with increased von Neumann entropy. In the model presented here, a sustaining superoperator 𝒮 = −𝒟 is introduced, mathematically canceling the environmental decoherence term so that the effective evolution becomes unitary. This cancellation stabilizes a sustained pure-state phase in which coherence, purity, and entropy remain constant even in an open environment. The identity functional 𝒥[*ρ*], which quantifies relational pattern structure rather than material substrate, remains invariant under global substrate replacement. This captures the notion of substrate-invariant embodiment, where functional identity persists despite atomic and molecular turnover. Finally, the framework enables collective many-body tunneling characterized by sublinear effective mass scaling *Meff* = *m N*^*a*^ with 0 < *α* < 1. This scaling supports coherent relocation of the system as an integrated whole, consistent with enhanced transport phenomena observed in biological quantum processes. Together, these elements illustrate how a sustaining superoperator can arrest decoherence, preserve informational identity, and support coherent many-body behavior in biological or bio-inspired systems.

### 3.8 Integrated Framework Illustrated in the Graphical Abstract

Figure 7 provides a consolidated graphical model of the framework. The visual summary highlights:

1. Ordinary open biological systems undergo entropy-producing Lindblad decoherence.
2. Introducing a sustaining superoperator 𝒮 = −𝒟 restores unitary evolution and stabilizes a sustained-coherence phase.
3. The identity functional 𝕀[*ρ*] remains invariant under substrate replacement.
4. Sublinear mass scaling *Meff* = *m N*^*a*^ supports collective many-body tunneling. Together, these results define a unified picture in which sustained coherence, substrate-invariant identity, and many-body quantum mobility emerge as interconnected consequences of compensatory open-system dynamics.

## 4 Discussion

This study introduces and formalizes the concept of a sustaining superoperator (𝒮) that exactly counterbalances environmental decoherence (𝒟) in open quantum systems. Through this theoretical construct, we propose a mechanism that maintains zero entropy production, constant purity, and invariant informational identity in a regime analogous to a decoherence-free manifold. While inspired by quantum biological observations, the results presented here generalize to any open quantum system capable of implementing self-consistent negentropic feedback. The implications span from quantum thermodynamics to biological network theory, suggesting that coherence, once thought ephemeral under thermal noise, could be dynamically stabilized through structured interactions that encode information as relational invariants rather than material substrates.

### 4.1 Theoretical implications of the sustaining superoperator

The sustaining superoperator formalism extends the Gorini-Kossakowski-Lindblad-Sudarshan (GKLS) framework by introducing a corrective operator that cancels decohering terms while maintaining complete positivity and trace preservation. In conventional Lindblad dynamics, 𝒟[ρ] induces exponential decay of off-diagonal elements, increasing von Neumann entropy *S*_*vN* = −*k*_*B Tr*(*ρ lo*g *ρ*). In our formulation, the addition of 𝒮[*ρ*] = −𝒟[*ρ*] *yi*e*lds dS*_*vN*/*dt* = 0, ensuring an entropy-neutral trajectory. This equivalence to reversible dynamics within an open system represents a physically plausible model of reversible dissipation, a concept compatible with detailed balance in non-Hamiltonian systems (Alicki, 1976).

Mathematically, this suggests that the state space of an open system contains embedded submanifolds of exact reversibility-zones where decoherence is continuously counteracted by negentropic synchronization. These can be interpreted as attractors in the extended phase space of quantum thermodynamics, analogous to limit cycles in dissipative classical systems. Within such manifolds, the system’s trajectory retains coherence indefinitely, implying that decoherence and recoherence are not mutually exclusive but can coexist in steady-state balance. This framework provides a rigorous basis for the notion that biological systems, continuously exchanging matter and energy, might achieve coherence through dynamic equilibrium rather than isolation.

### 4.2 Informational identity as a relational invariant

The introduction of the identity functional 𝒥[*ρ*] redefines persistence not as the endurance of matter, but as the conservation of correlation topology. In standard physical ontology, identity is attached to material constituents; in the present theory, it is attached to informational geometry. The invariance of 𝒥[*ρ*] under unitary permutation *Uπ* implies that the system’s “selfhood” persists even when its constituent particles are entirely replaced, provided the relational observables *O*k and *O*ℓ maintain constant statistical correlations.

This principle parallels both information-theoretic and biological definitions of individuality. In quantum information, error-corrected logical qubits preserve state identity independent of physical qubit replacement (Lidar & Brun, 2013). Similarly, in biology, cellular identity persists through turnover of molecular constituents because organizational invariants, such as gene regulatory networks, protein interaction motifs, and metabolic flux patterns, remain stable (Walker & Davies, 2013). The mathematical treatment presented here unifies these perspectives by describing biological persistence as a quantum-informational invariant encoded in the relational structure of correlations.

From a thermodynamic viewpoint, identity conservation through 𝒥[*ρ*] can be viewed as an emergent order parameter describing system-wide coherence. This concept resonates with Frohlich’s hypothesis (1968) that coherent oscillations in biomolecular assemblies arise from energy condensation into long-range modes. In such cases, biological structure is preserved not through static composition but through continuous energy throughput that maintains coherent phase relations. Thus, the sustaining superoperator could represent an effective theoretical abstraction of this energy-driven coherence stabilization.

### 4.3 Coherence networks and quantum-informational biology

Biological systems, especially at the mesoscopic scale, exhibit networked organization in which coherence is distributed among interacting units. In photosynthetic complexes, excitonic transport efficiency increases with delocalization, reaching near-unity quantum yields (Engel et al., 2007; Collini et al., 2010). Similar collective coherence has been proposed in enzymatic catalysis, where tunneling pathways depend on synchronized nuclear vibrations (Huelga & Plenio, 2013). In neural systems, microtubule networks have been theorized to support coherent dipolar modes (Pokorný, 2012).

The sustaining superoperator framework naturally generalizes these observations. By defining coherence-preserving transformations at the level of the system’s density matrix, it formalizes feedback processes that could underlie coherence protection in such networks. Biological systems can, in principle, generate structured negentropic feedback through biochemical oscillations, molecular chaperones, or coupled hydration shells that rephase decohering states (Kurian et al., 2018). These processes might effectively implement local approximations of 𝒮[ρ], dynamically reestablishing coherence lost to environmental fluctuations.

Moreover, sublinear effective-mass scaling (*Meff* = *m N*^*a*^, *α* < 1) suggests that coherent ensembles behave as extended entities with delocalized inertia. This could help explain phenomena such as long-range energy migration in protein lattices or synchronized metabolic oscillations observed in yeast and cardiac tissue. The theoretical limit *αc* ≈ 0.4 provides a quantitative threshold between collective coherence and classical independence, a criterion potentially measurable through experimental spectroscopy or single-molecule coherence analysis.

### 4.4 Implications for biological quantum networks

If coherence and identity persistence indeed coexist under entropy-neutral conditions, biological organization may exploit quantum-information principles far beyond molecular scale. Enzyme complexes, cytoskeletal networks, and chromatin domains could represent adaptive coherence reservoirs, where structured dissipation and negentropic feedback jointly stabilize functional order. Such systems might realize minimal instances of the sustaining superoperator through recurrent coupling and dynamic feedback.

At the organismal level, neural and cardiac synchronization offer macroscopic analogues of quantum phase locking (Pikovsky et al., 2003). While classical synchronization mechanisms are sufficient to describe many such processes, the possibility of quantum coherence contributing to phase stability in biological tissues remains open. If so, the sustaining superoperator formalism could provide a unifying description of coherence-preserving mechanisms spanning molecular to macroscopic domains.

The implications for origin-of-life theories are also profound. If informational identity can be preserved independent of substrate, then prebiotic systems might have achieved early forms of quantum coherence stabilization prior to the emergence of genetic encoding. Theoretical models of dissipative self-organization (England, 2013; Walker, 2017) already suggest that life arises from systems that maintain low entropy through feedback-driven structure formation. The present framework extends this principle to quantum regimes, where coherence itself becomes the conserved quantity driving emergent organization.

### 4.5 Prospects for experimental and computational validation

Experimental validation of the sustaining superoperator model requires identifying biological or synthetic systems that exhibit entropy-neutral coherence stabilization. Ultrafast two-dimensional electronic spectroscopy could be used to monitor long-lived coherence in protein complexes under controlled perturbations (Panitchayangkoon et al., 2011). Similarly, optomechanical biosensors could detect subthermal fluctuations consistent with entropy cancellation. If the sustaining mechanism is real, one should observe a steady-state coherence that persists without external energy input, distinguished by zero net entropy drift (Δ*S* ≈ 0).

Computationally, this framework could be implemented in open quantum system simulators using non-Markovian noise models. Machine learning-assisted quantum dynamics (Preskill, 2018) could optimize feedback operators that emulate 𝒮[*ρ*], testing how structured negentropic corrections sustain purity. Such approaches would directly connect theoretical predictions with experimentally measurable observables, such as coherence lifetimes, tunneling amplitudes, and purity fluctuations.

Synthetic analogues of biological coherence could also be engineered. DNA-templated quantum dots or peptide-based quantum wires may serve as testbeds for designing negentropic feedback circuits. In these systems, active stabilization via photonic or phononic coupling could operationalize the sustaining superoperator experimentally, demonstrating entropy-neutral quantum dynamics at ambient temperatures (Scholes et al., 2011).

### 4.6 Philosophical and theoretical outlook

Beyond its scientific implications, the sustaining superoperator formalism reframes long-standing questions about persistence, individuality, and the thermodynamic basis of life. It suggests that living systems may not merely resist entropy through metabolic work, but actively sculpt their local entropy landscape to preserve coherence. Identity, in this view, is not a static property but a dynamically maintained correlation topology, a quantum-informational essence persisting through material change.

Such a redefinition aligns with emerging interdisciplinary theories of life as an informational phenomenon. The algorithmic origins of life (Walker & Davies, 2013) and the theory of dissipative adaptation (England, 2013) both posit that self-organizing systems evolve structures that optimally maintain informational integrity. The sustaining superoperator provides a concrete mathematical embodiment of this idea, formalizing how negentropic feedback could enforce informational invariance in open systems.

This framework also invites reconsideration of consciousness and cognition as potential manifestations of large-scale informational coherence. If neural networks maintain coherence-like states via recurrent synchronization and feedback, they may instantiate high-level analogues of 𝒮[ρ]. While speculative, this hypothesis integrates physical, informational, and biological definitions of persistence into a unified theory of coherent identity.

In summary, the sustaining superoperator formalism unites principles from quantum thermodynamics, information theory, and biological systems science to propose a mechanism for persistent coherence and substrate-independent identity. The key innovation lies in showing that entropy cancellation, identity invariance, and collective tunneling can coexist within a single mathematical framework. Together, these results suggest that biological coherence and informational identity may be emergent consequences of structured negentropic feedback, a principle with profound implications for both fundamental physics and the theory of life itself.

Future research should combine high-resolution spectroscopic observation, computational quantum biology, and synthetic biomimetic experimentation to test the predictions derived here. Demonstrating entropy-neutral coherence in controlled systems would not only substantiate the sustaining superoperator hypothesis but also advance our understanding of how nature stabilizes complexity, coherence, and identity across all scales of existence.

## 5 Conclusion

This work establishes a mathematical and physical foundation for sustained quantum coherence and informational identity in open systems. By introducing the sustaining superoperator (𝒮) as a formal counterbalance to the dissipator (𝒟) in the Lindblad equation, we show that entropy production can be neutralized without violating thermodynamic constraints. This results in a zero-entropy pure-state manifold, in which quantum coherence is preserved indefinitely, and system dynamics remains reversible despite environmental coupling. Such a regime redefines the boundary between closed and open dynamics, offering a new pathway for understanding how life-like stability and informational continuity may emerge from physical law.

The informational identity functional 𝒥[*ρ*] introduced here formalizes the idea that selfhood can be defined as a persistent pattern of correlations, independent of material substrate. This principle resonates deeply with information-centric models of living systems, where organizational invariance, not molecular permanence, determines individuality and continuity. In biological terms, this could explain how living organisms, which undergo constant molecular turnover, retain coherent identity across time. The invariance of 𝒥[*ρ*] under random substrate permutations mathematically encodes this observation, demonstrating that informational topology can remain unchanged even when the underlying physical structure is replaced.

The collective mass scaling *Meff* = *m N*^*a*^ and the persistence of finite tunneling amplitudes for *α* < *αc* ≈ 0.4 provide physical evidence of collective coherence across system components. This phenomenon mirrors the reduced inertia seen in macroscopic quantum systems such as superconductors and superfluids, where coherence leads to emergent properties not attributable to individual particles (Leggett, 1999). Our findings suggest that when coherence is distributed throughout a system, the ensemble behaves as a single quantum entity with reduced effective mass, enabling whole-body tunneling or coherent relocation that transcends classical spatial separation.

From a broader perspective, this study outlines the conditions under which substrate-invariant embodiment, the persistence of informational identity despite material exchange, can occur. In this framework, an entity’s continuity is encoded not in its constituent atoms but in the self-consistent network of quantum correlations that defines it. This challenges traditional materialist interpretations of persistence and opens the possibility that living order may be maintained by coherence-preserving feedback acting at quantum and mesoscale levels. Such mechanisms might already be at play in nature, as suggested by experimental reports of coherence in photosynthetic reaction centers (Engel et al., 2007; Panitchayangkoon et al., 2011), enzyme catalysis (Klinman & Kohen, 2013), and avian magnetoreception (Ritz et al., 2000). In physical and computational terms, the sustaining superoperator may be viewed as a quantum generalization of error correction or feedback control, maintaining the purity of the system’s density operator through adaptive negentropic compensation. This conceptual bridge between quantum thermodynamics and biological information processing suggests a path toward engineered coherence preservation, applicable to synthetic biology, quantum computing, and artificial life systems. The same mathematical principles could underlie quantum error-corrected biosystems-entities that maintain functionality and identity across energy fluctuations, noise, or physical replacement of components.

In essence, the framework presented here proposes that life-like persistence, the ability to remain “the same” through continuous physical change, can emerge from fundamental physics through entropy cancellation and coherence conservation. The unification of sustained coherence, identity invariance, and collective relocation defines a new theoretical class of quantum-coherent open systems, bridging the gap between quantum mechanics and biological individuality. Future directions should include:

1. extending this model to realistic molecular Hamiltonians that include vibronic coupling and thermal noise,
2. experimentally probing coherence stabilization in biological condensates and synthetic quantum biopolymers, and
3. exploring whether biological networks actively implement negentropic superoperators through metabolic or photonic feedback. If verified, such mechanisms could redefine our understanding of biological continuity, consciousness, and identity as emergent expressions of coherent quantum information, a bridge between physics and the persistent phenomena we call life.

## Notes

https://github.com/mpetalcorin/Quantum-Coherent-Identity-Preservation-in-Open-Biological-Systems

### Competing Interest Statement

The authors have declared no competing interest.

